# Comammox bacterial preference for urea influences its interactions with aerobic nitrifiers

**DOI:** 10.1101/2023.07.11.548560

**Authors:** Katherine Vilardi, Juliet Johnston, Zihan Dai, Irmarie Cotto, Erin Tuttle, Arianna Patterson, Aron Stubbins, Kelsey Pieper, Ameet Pinto

## Abstract

While the co-existence of comammox bacteria with canonical nitrifiers is well documented in diverse ecosystems, there is still a dearth of knowledge about the mechanisms underpinning their interactions. Understanding these interaction mechanisms is important as they may play a critical role in governing nitrogen biotransformation in natural and engineered ecosystems. In this study, we tested the ability of two environmentally relevant factors (nitrogen source and availability) to shape interactions between strict ammonia and nitrite-oxidizing bacteria and comammox bacteria in continuous flow column reactors. The composition of inorganic nitrogen species in reactors fed either ammonia or urea was similar during the lowest nitrogen loading condition (1 mg-N/L), but higher loadings (2 and 4 mg-N/L) promoted significant differences in nitrogen species composition and nitrifier abundances. The abundance and diversity of comammox bacteria were dependent on both nitrogen source and loading conditions as multiple comammox bacterial populations were preferentially enriched in the urea-fed system. In contrast, their abundance was reduced in response to higher nitrogen loadings in the ammonia-fed system likely due to ammonia-based inhibition. The preferential enrichment of comammox bacteria in the urea-fed system could be associated with their ureolytic activity calibrated to their ammonia oxidation rates thus minimizing ammonia accumulation to inhibitory levels. However, an increased abundance of comammox bacteria was not associated with a reduced abundance of nitrite oxidizers in the urea-fed system while a negative correlation was found between them in the ammonia-fed system; the latter dynamic likely emerging from reduced availability of nitrite to strict nitrite oxidizers at low ammonia loading conditions.

**Importance:** Nitrification is an essential biological process in drinking water and wastewater treatment systems for managing nitrogen and protecting downstream water quality. The discovery of comammox bacteria and their detection alongside canonical nitrifiers in these engineered ecosystems has made it necessary to understand the environmental conditions that regulate their abundance and activity relative to other better-studied nitrifiers. This study aimed to evaluate two important factors that could potentially influence the behavior of nitrifying bacteria, and therefore impact nitrification processes. Colum reactors fed with either ammonia or urea were systematically monitored to capture changes in nitrogen biotransformation and the nitrifying community as a function of influent nitrogen concentration, nitrogen source, and reactor depth. Our findings show that comammox bacterial abundance decreased and that of nitrite oxidizers increased with increased ammonia availability, while their abundance and diversity increased with increasing urea availability without driving a reduction in the abundance of canonical nitrifiers.

## Introduction

Comammox bacteria are routinely detected alongside strict ammonia oxidizing bacteria (AOB) and nitrite oxidizing bacteria (NOB) in both drinking water and wastewater systems (Cotto et al., 2020; Fowler et al., 2018; Pinto et al., 2015; Poghosyan et al., 2020; Roots et al., 2019; K. J. Vilardi et al., 2022; Wang et al., 2017; Yang et al., 2020; Zheng et al., 2023) but insights into the factors influencing their abundance, activity, and interactions in these environments are still limited. Interactions between AOB and NOB have been extensively studied including the impact of process and environmental conditions such as oxygen supply, ammonia concentration, and temperature (Pérez et al., 2014; Seuntjens et al., 2018; Sliekers et al., 2005). However, the presence of comammox bacteria within these communities requires a re-evaluation of these interactions and the collective response of nitrifying consortia to changes in environmental and/or process conditions. Our understanding of the role and ecological niche of comammox bacteria within complex nitrifying communities is further restricted by limited physiological insights due to the existence of only a few cultured representatives and/or enrichments, all belonging to clade A1 (Daims et al., 2015; Ghimire-Kafle et al., 2023; Sakoula et al., 2021).

Ammonia availability is likely an important factor governing interactions between strict AOB and comammox bacteria. For instance, comammox bacterial cultures and enrichments have shown significantly higher affinity for ammonia compared to strict AOB (Ghimire-Kafle et al., 2023; Kits et al., 2017; Sakoula et al., 2021). Thus, comammox bacteria may outcompete strict AOB in ammonia-limited environments such as drinking water systems. Further, different comammox bacteria may exhibit varying preferences for ammonia concentration ranges and these may be dictated not just by ammonia affinities, but also by potential inhibition at higher concentrations. For example, ammonia oxidation by *Ca.* Nitrospira krefti was partially inhibited at relatively low ammonia concentrations (25 μM) (Sakoula 2021) which was not observed for *Ca.* Nitrospira inopinata (Kits 2017). Comammox bacteria may also exhibit clade/sub-clade dependent preferences for ammonia availability and/or environments. For example, clade A1 comammox bacteria associated with *Ca.* Nitrospira nitrosa are typically found at higher abundances than canonical nitrifiers in some wastewater systems (Cotto 2020, Wang 2018, Xia 2018, Zheng 2023) and sometimes as the principal aerobic ammonia oxidizers in a wastewater system (Vilardi and Cotto 2023). In the latter situation, ammonia oxidation dominated by comammox bacteria could also adversely impact *Nitrospira*-NOB by limiting nitrite availability through complete nitrification to nitrate at low ammonia concentrations; however, the relationship between the two *Nitrospira* groups is not well understood.

Nitrogen source could also have a significant effect on interactions between nitrifiers. For instance, ureolytic activity may enable access to ammonia derived from urea in engineered systems (e.g., wastewater treatment), as well as natural systems such as freshwater ecosystems (Solomon et al., 2010). Genes for urea degradation accompanied by a diverse set of urea transporters are ubiquitously found in genomes of all comammox bacteria (Palomo et al., 2018). Their ability to grow in urea is supported by the enrichment of multiple species of comammox bacteria urine-fed membrane bioreactors (J. Li et al., 2021) and enrichment of comammox bacteria when supplied with urea (J. Li et al., 2021; Zhao et al., 2021). Some *Nitrospira*-NOB are also capable of catalyzing ammonia production through urea degradation and thus, potentially regulating nitrite availability via cross feeding of ammonia to strict AOB (Koch et al., 2015); this could potentially influence competition between canonical nitrifiers and comammox bacteria.

The present work aimed to investigate comammox bacterial preferences and potential interactions with canonical nitrifiers subject to different nitrogen sources and loadings. We operated two continuous-flow laboratory-scale column reactors with granular activated carbon (GAC) containing all three nitrifying groups and supplied the reactors with either ammonia or urea at three different influent nitrogen loadings. Our goal was to infer (1) nitrogen source (i.e., ammonia and urea), species (i.e., urea, ammonia, nitrite), and concentration preferences of nitrifying groups and (2) their potential interactions by quantitatively measuring their differential sorting within column reactors over time in the context of their genome-resolved metabolic capabilities.

## Materials and methods

### Reactor operation

Two laboratory-scale column reactors (diameter = 1”, height = 10”) were packed with GAC (packed height = 3’’) and operated with an approximately 1” of water head above the GAC to ensure the media was fully saturated. The systems were each packed with 35 g of GAC from the City of Ann Arbor, Michigan Drinking Water Treatment Plant (DWTP). The two reactors were fed with synthetic groundwater media (Smith et al., 2002). Stock solution for the inorganic compounds in the media was prepared with 3.88 g/L MgCl_2_, 2.81 g/L CaCl_2_, 13.68 g/L NaCl, 6.90 g/L K_2_CO_3_, 17.75 g/L Na_2_SO_4_, and 0.88 g/L KH_2_PO_4_. The organic compound stock solution contained 3.75 g/L of glucose (C_6_H_12_O_8_) and a third sodium bicarbonate solution was prepared with 30 g/L of NaHCO_3_. Influent media was then prepared in 10-L autoclaved carboys with 1 mL/L of the inorganic and organic compound stock solutions and 10 mL/L of the sodium bicarbonate stock solution. The two reactors were fed influent amended with stock solutions of ammonium chloride (NH_4_Cl) or urea (CH_4_N_2_O). Influent media was pumped at 1.15 L/day with the peristaltic pump resulting in an empty bed contact time (EBCT) of approximately 48 minutes. Both reactors were fed influent at three different nitrogen concentrations over the experimental period. Columns reactors were maintained in conditions 1 (1 mg-N/L) and 2 (2 mg-N/L) for eight weeks and in condition 3 (4 mg-N/L) for six weeks.

### Sample collection and processing

Influent and effluent were sampled twice weekly, while five samples spaced approximately 0.5” apart along the depth of the GAC column were collected weekly to capture depth-wise nitrogen species concentrations. The five sections are defined as sections 0.5 (top), 1, 1.5, 2 and 2.5 (bottom). All aqueous samples were filtered through 0.22 μm filters (Sartorius Minisart NML Syringe Filter - Fisher Scientific 14555269). GAC media samples were collected at week 0 followed by weeks six, seven and eight for conditions 1 and 2, and week six for condition 3. GAC media samples (0.3 grams) were collected from three locations along the reactor bed: one within the top 0.5” of the reactor, one mid-filter depth (1.5”), and another at the bottom approximately 3” from the top of the reactor location and were stored for DNA extraction in Lysing Matrix E Tubes (MP Biomedical - Fisher Scientific MP116914100). After each sampling event, the amount of GAC taken was replaced with virgin GAC which was mixed with the remaining GAC by first fluidizing the filter media with 50 mL deionized water followed by backwashing with air for 5 minutes.

### Chemical Analysis

Hach TNT Vials were used to determine concentrations of ammonia (TNT832), nitrite (TNT839), nitrate (TNT835), and total alkalinity (TNT870). All samples were analyzed on a Hach DR1900 photospectrometer (Hach—DR1900-01H). Influent and effluent pH was determined using a portable pH meter (Thermo Scientific™ Orion Star™ A221 Portable pH Meter – Fisher Scientific 13-645-522). A Shimadzu TOC-L (total organic carbon analyzer) with a TNM-L attachment (total nitrogen unit) (Stubbins and Dittmar 2012) was used to measure total dissolved nitrogen in influent and effluent samples using certified DOC/TDN standards (deep seawater reference (DSR): low carbon seawater, LSW, deep seawater reference material) (Batch 21 Lot 11-21, 1). Urea concentrations in samples collected from the urea-fed reactors were determined by subtracting the total inorganic nitrogen measured (i.e., sum of ammonia, nitrite, and nitrate) in each sample from influent urea concentration.

### Nitrogen biotransformation rate calculations

Rates of nitrogen biotransformations were calculated from the concentration profiles of ammonia, NO_x_ (nitrite plus nitrate), nitrate, and total inorganic nitrogen (sum of ammonia, nitrite, and nitrate) measured along the column reactor depths. Rates were calculated for six sections of the columns: 0-0.5, 0.5-1, 1-1.5, 1.5-2, 2-2.5, and section 2.5-3 inches. Volumetric rates (mg-N/L packed GAC/h) were obtained by multiplying the concentration differences between the profile layers by the influent flow rate and dividing by the volume of packed GAC between the profile layers (V=6.4 mL packed GAC in each layer).

### DNA extraction and qPCR assays

DNA was extracted from all GAC samples (n=43) which included the inocula and samples collected at all time points and locations during each condition. Extractions were performed using Qiagen’s DNeasy Powersoil Pro (Qiagen, Inc – Cat. No. 47014) with a few modifications. GAC in lysing matrix tubes with 800 μL of CD1 was vortexed briefly and placed in a 65°C water bath for 10 minutes. After heating, 500 μL of phenol:chloroform:isoamyl Alcohol (25:24:1, v/v) (Invitrogen™ UltraPure™ - Fisher Scientific 15-593-031) was added to the lysing tube and bead beating proceeded with four 40 second rounds on the FastPrep-24 instrument (MP Biomedical – Cat. No. 116005500) with lysing tubes placed on ice for two minutes between rounds. Samples were then centrifuged for one minute and 600 μL of the aqueous phase was used for DNA extractions on the Qiacube (Qiagen, Inc—Cat No. 9002160) protocol for Powersoil Pro. A reagent blank was included in each round of extractions as a negative control. DNA concentrations were measured using a Qubit with the dsDNA Broad Range Assay (Invitrogen™ - Fisher Scientific Q32850). Extracted DNA was stored at -80°C until further processing.

qPCR assays were conducted using Applied Biosystems 7500 Fast Real-Time PCR instrument. Primer sets listed in supplemental table 1 were used to target the 16S rRNA gene of AOB (Hermansson and Lindgren 2001), 16S rRNA gene of *Nitrospira* (Graham 2007), *amoB* gene of clade A comammox bacteria (Vilardi 2022), and 16S rRNA gene of total bacteria (Caporaso 2011). The qPCR reactions were performed in 20 μL volumes, which contained 10 μL Luna Universal qPCR mastermix (New England Biolabs Inc., Fisher Scientific Cat. No. NC1276266), 5 μL of 10-fold diluted template DNA, primers at concentrations listed in supplemental table 1 and DNA/RNAase free water (Fisher Scientific, Cat. No. 10977015) to make the remaining volume. Each sample was subjected to qPCR in triplicate. The cycling conditions consisted of initial denaturing at 95°C for 1 minute, 40 cycles of denaturing 95°C for 15 seconds, annealing times and temperatures listed in supplemental table 1, and extension at 72°C for 1 minute. Three different sets of gBlock standards (Integrated DNA Technology gBlocks® Gene Fragments 125-500 bp) targeting the 16S rRNA gene of total bacteria and *Nitrospira*, 16S rRNA gene of AOB and *amoB* gene of clade A comammox bacteria were used to establish a 7-point standard curve for each respective assay (supplemental table 2). The qPCR efficiencies for all assays are listed in supplemental table 1.

### 16S rRNA gene amplicon sequencing and data analysis

DNA extracts (triplicate per sample) from all samples were submitted for sequencing of the V4 hypervariable region of the 16S rRNA gene at the Georgia Institute of Technology Sequencing Core. The MiSeq v2 kit was used to generate 250 bp pair-end reads using the 515F (Parada et al., 2016) and 806R (Apprill et al., 2015) primers with overhang of Illumina adapters. Removal of adapter and primer sequences from the resultant sequencing data was carried out using cutadapt v4.2. Amplicon sequencing data processing and quality filtering was performed using DADA2 v1.22.0 (Callahan 2016) in R v4.1.2. to infer amplicon sequencing variants (ASVs) using the pipeline for paired-end Illumina MiSeq data. The SILVA nr v.138.1 database was used for taxonomic assignment of ASVs with a minimum bootstrap confidence threshold of 80. The ASV table was rarefied with ‘rarefy_even_depth’ function from the R package phyloseq v1.38.0 to the sample with the smallest library size. The relative abundance of ASVs in each sample were calculated by dividing ASV reads counts in the sample by the total number of samples read counts.

### Metagenomic sequencing, assembly, and binning

DNA extracted from samples taken at week six from the top layer of the ammonia- and urea-fed reactors during condition 3 were submitted for sequencing on the Illumina NovaSeq platform with a SP flow cell at the Georgia Institute of Technology Sequencing Core. Similar workflows and tools utilized in Vilardi 2022 and Cotto 2023 (Cotto et al., 2023; K. J. Vilardi et al., 2022) were applied here to assemble and characterize metagenome assembled genomes (MAGs). Briefly, raw paired-end reads were quality filtered using fastp (Chen et al., 2018) and further mapped to the Univec database to remove contaminated reads. Samtools (Danecek et al., 2021) was used to sort the resulting bam files and bedtools (Quinlan & Hall, 2010) was used to convert them to fastq files. Assemblies were generated for each sample separately using metaSpades (Nurk et al., 2017) with kmer sizes 21, 33, 55 and 77. The two resulting fasta assemblies were indexed with bwa index and filtered pair-ended reads were mapped to back to their respective assemblies with bwa mem (H. Li & Durbin, 2009). The subsequent sam files were converted to bam files using appropriate samtools flags to retain only mapped reads.

Metabat2 (Kang et al., 2019) was used to bin contigs longer than 2000 bp followed by CheckM (Parks et al., 2015) to determine completeness and contamination levels of metagenome assembled genomes (MAGs) which were then classified using the Genome Taxonomy Database Tool Kit with database release 207 (Chaumeil et al., 2019; Parks et al., 2018). Open reading frames of coding regions predicted using prodigal (Hyatt et al., 2010) were annotated against the KEGG database (Kanehisa et al., 2016) with kofamscan (Aramaki et al., 2020). Up-to-date Bacterial Core Gene pipeline (Na et al., 2018) was used to construct maximum likelihood trees using a set 92 extracted and aligned single copy genes from assembled *Nitrospira*-like and *Nitrosomonas-*like MAGs and references. A set of dereplicated MAGs was generated from MAGs recovered from the ammonia- and urea-fed systems at an ANI threshold of 99% using dRep (Olm et al., 2017). Reads from the ammonia- and urea-fed samples were mapped to the set of dereplicated MAGs to calculate breadth of coverage (i.e., percent of genome covered by reads) and relative abundance using coverM (https://github.com/wwood/CoverM).

16S rRNA gene sequences of a minimum length of 500bp were reconstructed from the metagenomes of both samples using MATAM v1.6.1 (Pericard et al., 2018) to establish the linkage between ASVs generated from 16S rRNA gene amplicon sequencing and MAGs associated with nitrifying bacteria. To achieve a more comprehensive reconstruction, recursive random subsampling of different depths, i.e., 1%, 5%, 10%, 25%, 50%, 75% and 100%, was performed, followed by dereplication at 99.9% identity using USEARCH v11.0.667 (Edgar, 2010; Song et al., 2022). Only the longest sequence from each cluster was retained for downstream analysis. Furthermore, ASVs were aligned against the MATAM recovered 16S rRNA gene sequences and extracted 16S rRNA genes from MAGs by Barrnap using BLASTn v2.13.0 (Camacho et al., 2009), and only ASV hits of 100% identity and 100% coverage were considered as linkage candidates between ASV and MAG unless the alignment interrupted at the end of the reference sequence.

### Statistical Analysis

All statistical analysis was performed in R v4.1.2 (RCore team., 2021). A significant difference in effluent concentrations of ammonia, nitrite, and nitrate in the ammonia- and urea-fed was tested using the non-parametric Wilcox rank sum test. Ratios of effluent NO_x_ to influent nitrogen concentrations as a proxy for ammonia consumption in both systems was compared with the Student’s T-test for conditions 2 and 3 while condition 1 required the systems be compared with the Welch’s T-test due to unequal variance between the ammonia- and urea-fed systems. The data distribution and variance for all tests was checked with the Shapiro-Wilks and Levene test, respectively. We tested if the microbial community in GAC samples clustered significantly by nitrogen source, loading condition, and reactor depth using the Bray-Curtis dissimilarities calculated from the ASV abundance table and applying a PERMANOVA test using the adonis function in the R package vegan v2.6-4 (*Vegan: Community Ecology Package*, 2022). Correlations between microbial community composition and concentrations of ammonia, nitrite, and nitrate measured in the biofiltrations were calculated with the Mantel test. The mean relative abundances of nitrifier ASVs and qPCR-based relative abundances of comammox bacteria, strict AOB and *Nitrospira*-NOB were compared in both systems across the nitrogen loading conditions using ANOVA. Pearson correlation was to statistically quantify and test the significance of the relationship between the abundance of comammox bacteria and Nitrospira-NOB in both systems.

### Data availability

Raw fastq files for amplicon sequencing and metagenomic sequencing data, metagenomic assembly, and curated MAGs are available via NCBI bioproject submission number SUB13674413.

## Results

### Nitrogen biotransformation in ammonia and urea fed systems

Urea and ammonia fed systems had similar concentrations of ammonia, nitrite, and nitrate in the effluent (Wilcoxon p>0.05) (Supplemental Figure 1) and similar depth-wise distributions of inorganic nitrogen species (Figure 1A) at lowest influent loading condition (condition 1 - 1 mg-N/L). Majority of the influent nitrogen (∼70%) was complete oxidized to nitrate in the topmost portion of the reactors (section In-0.5) (Figure 1A and 1B). Rates of ammonia oxidation (4.99 (±1.95) mg-N/L packed GAC/h) and urea degradation (5.13 (± 0.70) mg-N/L packed GAC/h) were highest in section In-0.5 and were nearly equal to the rate of nitrate production (5.31 (± 1.55) and 4.79 (± 0.83) mg-N/L packed GAC/h, respectively) (Figure 1B). Increasing the influent nitrogen concentrations to 2 mg-N/L (i.e., condition 2) led to significantly higher nitrite accumulation in the ammonia-fed compared to the urea-fed reactor (Wilcoxon p < 0.05) due to an imbalance between ammonia oxidation (10.91 (± 1.49) mg-N/L packed GAC/h) and nitrate production (8.51 (± 2.03) mg-N/L packed GAC/h) rates in section In-0.5 of the ammonia-fed system (Figure 1B). In contrast, average urea degradation rate (8.25 (± 2.21) mg-N/L packed GAC/h) was nearly equal to the nitrate production rate (6.85 (± 1.67) mg-N/L packed GAC/h) in the top section of the urea-fed reactor. The nitrite accumulation was exacerbated at the higher influent nitrogen loading condition (condition 3: 4 mg-N/L) with significantly higher effluent nitrite concentrations in the ammonia-fed (1.00 (±) mg-N/L) compared to the urea-fed system (0.10 (± 0.04) mg-N/L). The average ammonia oxidation rate (15.02 (± 2.67) mg-N/L packed GAC/h) in the top section of the ammonia-fed system was 1.8 times higher than the nitrate production rate (8.50 (± 2.03) mg-N/L packed GAC/h). The rates of nitrate production in section In-0.5 were similar between conditions 2 and 3 (8.56 (± 1.47) and 8.50 (± 2.03) mg-N/L packed GAC/h, respectively) in the ammonia-fed system indicating maximum rates of nitrate production had been reached. Interestingly, ammonia accumulation in the urea-fed systems resulted in similar effluent ammonia concentrations as the ammonia-fed systems for conditions 1 and 2 suggesting that both reactors had reached their ammonia oxidation capacity across the entire depth of the reactors.

**Figure 1:**
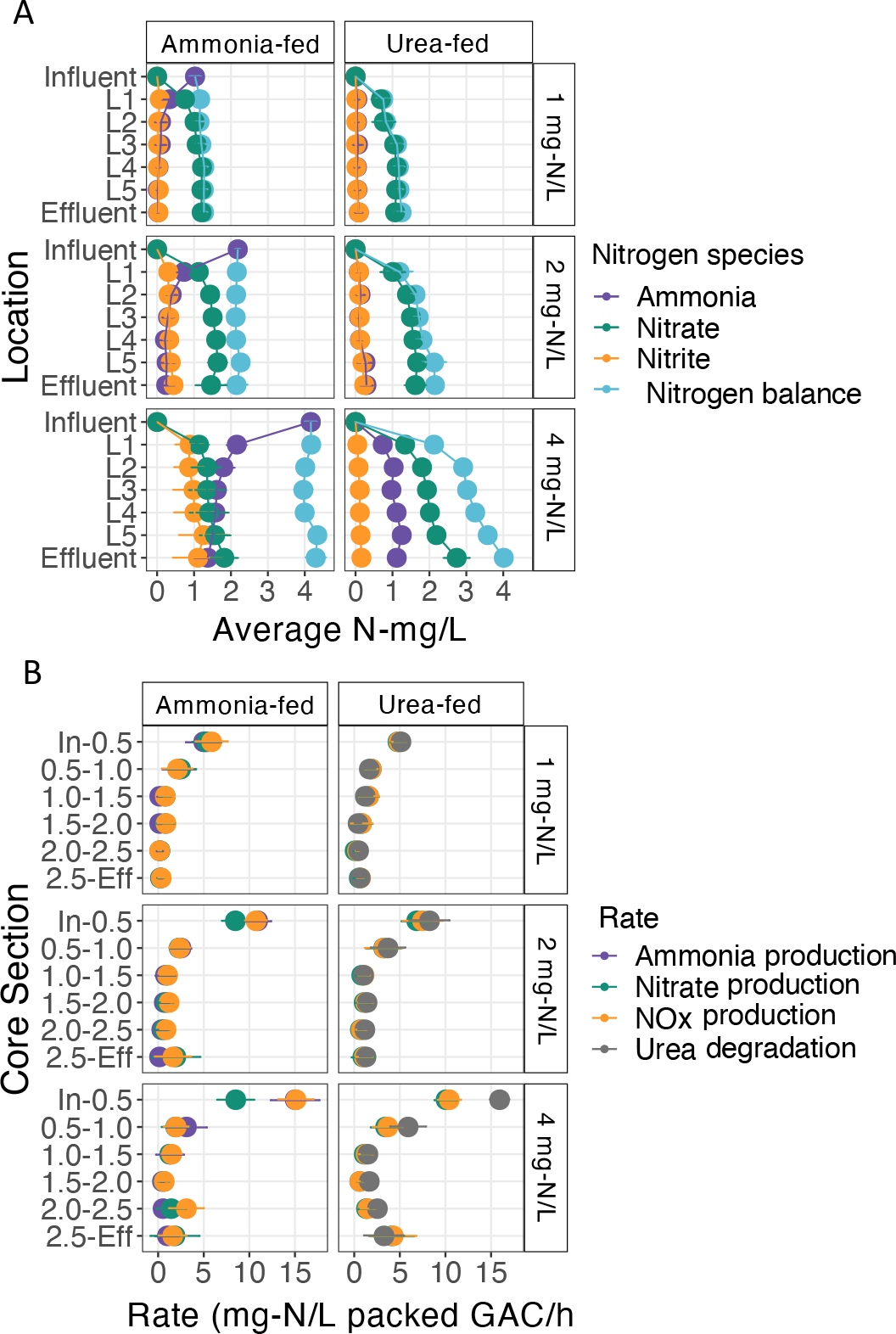
(A) Concentrations of ammonia, nitrite, and nitrate measured at five depths along the reactors. Measurements were taken at 0.5”, 1.0”, 1.5”, 2.0”, and 2.5” from the top of GAC in the reactors. Data points represent the average concentration obtained from the different reactor depths along with error bars for standard deviation. (B) Rates of nitrogen biotransformation along the depths of the reactors. Rates are colored by the type of nitrogen biotransformation (purple = ammonia oxidation, green = nitrate production, orange = NOx production, and grey = urea degradation). Data points represent the average rates obtained from the different reactor depths along with error bars for standard deviation.

### GAC microbial community composition

Rarefaction to the smallest library size (n=70354 reads) resulted in retention of 1738 ASVs out of 2400 constructed from the V4 hypervariable region of the 16S rRNA gene. ASVs with the highest relative abundance belonged to the class *Gammaproteobacteria* (5.09 (± 1.96) % - ASV 1 and 4.62 (± 2.96) % - ASV 2), *Vicinamibacteria* (3.70 (± 1.49) % - ASV 3), *Nitrospiria* (3.50 (± 1.63) % - ASV 4 and 2.72 (± 1.40) % - ASV 6), and *Alphaproteobacteria* (3.40 (± 1.56) % - ASV 5). Microbial community composition was shaped significantly by nitrogen loading condition (ANOSIM R=0.628, p < 0.05), nitrogen source (ANOSIM R = 0.134, p < 0.05), and reactor section (i.e., GAC sampling point, top (L1), middle (L3), bottom (L5)) (ANOSIM R = 0.163, p < 0.05) (Supplementary Figure 2A). Nitrogen source (i.e., ammonia-fed vs urea-fed) played a more significant role in shaping the overall microbial community for condition 2 (PERMANOVA R = 0.224, p < 0.05) as compared to condition 1 (PERMANOVA R = 0.104, p > 0.05) or condition 3 (PERMANOVA R = 0.412, p > 0.05). Community composition of the two reactors may not exhibit a strong difference for condition 1 due to very similar depth-wise nitrogen species profiles (Figure 1A and 1B), while differences during condition 3 may not be flagged as significant due to the limited data points. In the ammonia-fed system (Figure 2A), the microbial community separated into distinct clusters based on nitrogen loading condition (ANOSIM R =0.614, p < 0.05) but not by reactor depth (ANOSIM R = 0.077, p > 0.05). In contrast, both nitrogen loading (ANOSIM R = 0.665, p < 0.05) and reactor depth (ANOSIM R = 0.204, p < 0.05) were significantly associated with differences in microbial community composition for the urea-fed system (Figure 2B).

**Figure 2:**
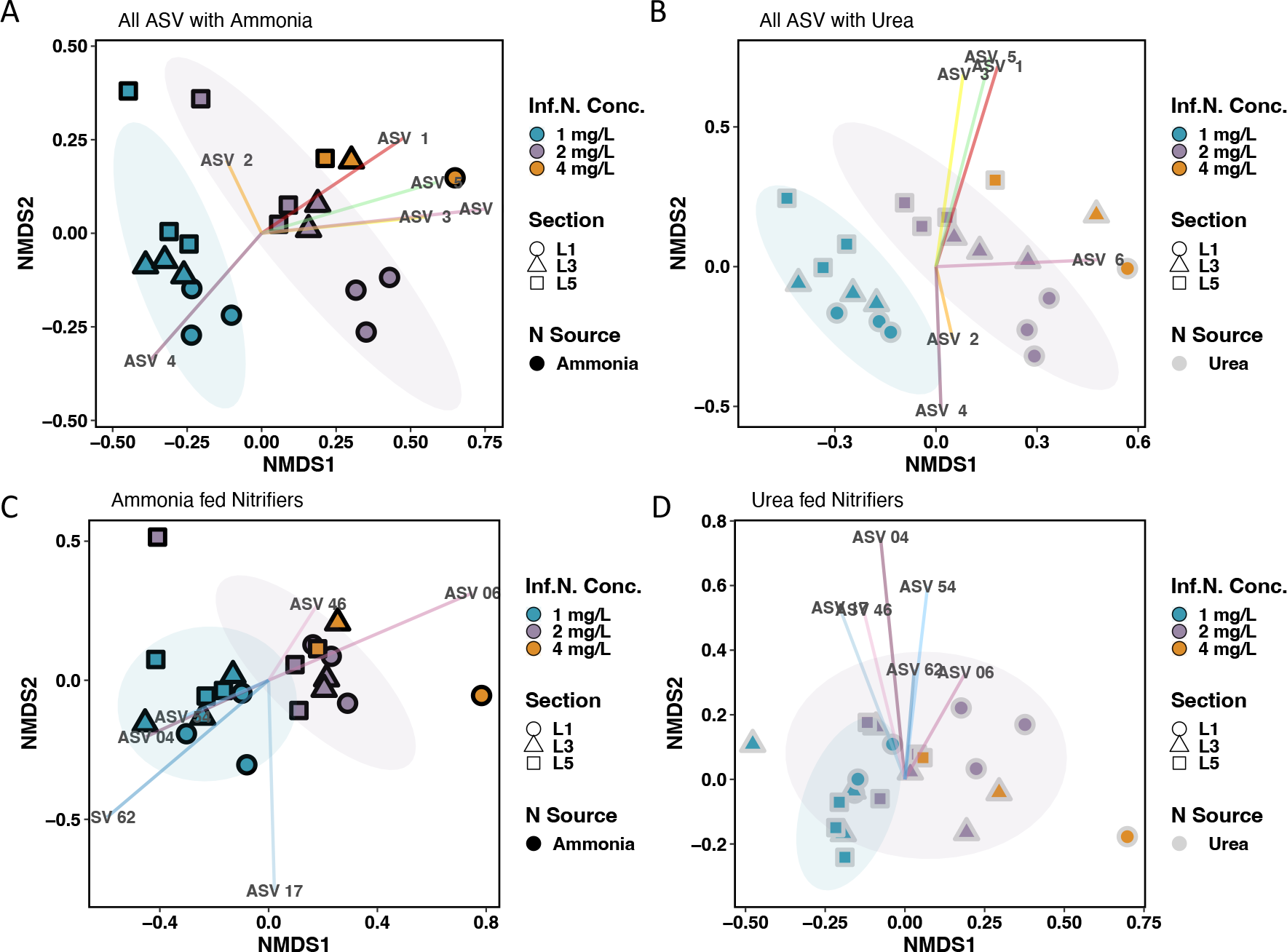
NMDS plots constructed with the abundance tables of all ASVs in the (A) ammonia-fed system, and (B) urea-fed system and nitrifier ASVs in (C) the ammonia-fed and (D) in the urea-fed system (D). Blue-, purple- and orange-colored points are GAC samples collected during conditions 1, 2, and 4, respectively. Shape symbolizes the reactor depth the GAC samples were taken from (L1 = top (circle), L3 = middle (triangle), and L5 = bottom (square)). The outline color of shapes represents the system the GAC was collected from (gray = urea-fed, black = ammonia-fed).

Compositional differences in nitrifier communities were evaluated with ASVs classified as *Nitrospira*- (9 ASVs) and *Nitrosomonas*-like (5 ASVs) bacteria; this is in line with our previous work which found nitrifiers belonged to only these genera in GAC samples from the same biofiltration system (Vilardi 2022). Nitrifier communities in the ammonia- and urea-fed systems were significantly dissimilar during conditions 1 (PERMANOVA R = 0.319, p < 0.05) and 2 (PERMANOVA R = 0.368, p < 0.05) (Supplementary Figure 2B). Nitrogen source explained a greater variance between communities in condition 3 (PERMANOVA R = 0.406) but was found to be insignificant potentially due to fewer data points. In both the ammonia- and urea-fed systems, nitrifier community composition was most dissimilar between conditions 1 and 4 (PERMANOVA R = 0.607 and 0.536, p < 0.05) (Figure 2C, D). Collectively, our results show that composition of both the whole community and nitrifiers were significantly shaped by nitrogen source and availability. The largest impacts were consistently observed when comparing the lowest and highest nitrogen loadings conditions.

### Impact of nitrogen source and loading conditions on nitrifying bacteria

In the ammonia and urea-fed systems, four *Nitrospira*-like (ASVs 4, 6, 46, 236) and three *Nitrosomonas-*like ASVs (ASVs 17, 54, and 62) were detected. Mapping ASVs 4, 6, and 46 against *Nitrospira* reference genomes and full length 16S rRNA sequences indicated ASV 4 was a *Nitrospira lenta*-like strict NOB (100% ID to NCBI accession number KF724505) and ASV 6 and 46 were *Nitrospira nitrosa*-like comammox bacteria (both 100 % ID to NZ_CZQA00000000). All *Nitrosomonas*-like ASVs shared high sequence similarity (>98% ID) with 16S rRNA gene sequences within *Nitrosomonas* cluster 6a (*Nitrosomonas ureae* (NZ_FOFX01000070) and Is79 (NC_015731).

To further evaluate the classification of the *Nitrospira* ASVs, 16S rRNA gene sequences were reconstructed using short reads obtained from metagenomic sequencing of samples taken from the top of the ammonia- and urea-fed reactors. ASVs 6, 46, and 4 uniquely mapped with 100% sequence identity to one, four, and four reconstructed 16S rRNA gene sequences, respectively. Phylogenetic analyses clustered all sequences corresponding to ASVs 6 and 46 with comammox bacterial species *Ca.* Nitrospira nitrificans while all ASV 4 matches clustered separately with *Nitrospira lenta* (Figure 3A). This further supports that two of the dominant Nitrospira ASVs (6 and 46) belonged to comammox bacteria and ASV 4 was strict *Nitrospira*-NOB. Dominant *Nitrosomonas*-like ASVs 17, 54, and 62 uniquely mapped with 100% sequence identity to five, four, and three reconstructed 16S rRNA gene sequences, respectively. Phylogenetic placement of the matches revealed those associated with ASV 17 clustered with *Nitrosomonas ureae* and sp. AL212 which suggests ASV 17 belongs to a lineage of urease-positive strict AOB (Figure 3B). ASV 62 matches were a part of the same main branch but formed a separate cluster with Nitrosomonas Is79A3 and sp. Nm86. All ASV 54 matches clustered with uncultured *Nitrosomonas*.

**Figure 3:**
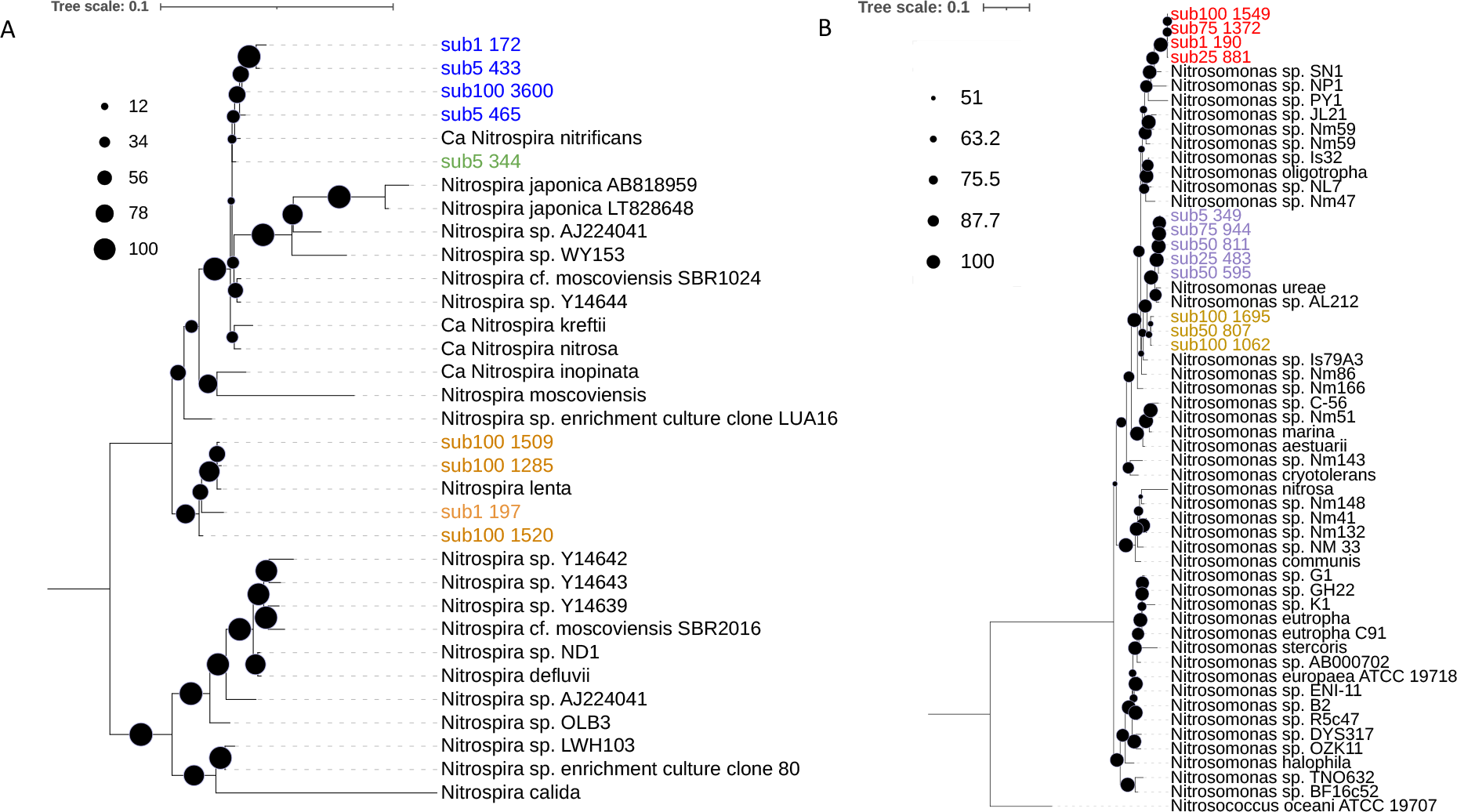
Maximum likelihood tree for 16S rRNA gene sequences for (A) Nitrospira and (B) Nitrosomonas. Color denotes the ASV matches: ASV 46 (Blue), ASV 6 (Green), ASV 4 (Orange), ASV 54 (Red), ASV 17 (Purple), and ASV 62 (Yellow).

In the ammonia-fed system, the average relative abundance of *Nitrospira lenta*-like ASV 4 increased with increasing nitrogen loadings (1 mg-N/L: 2.63 (±0.45) %, 2 mg-N/L: 3.89 (± 1.23) %, 4 mg-N/L: 5.40 (± 1.75) %) with its abundance significantly higher in condition 3 compared to condition 1 (ANOVA/Tukey, p < 0.05) (Supplementary Figure 3). In contrast, the average relative abundance of comammox*-*like ASV 6 decreased with increased nitrogen loadings (1 mg-N/L: 2.67 (± 0.70)%, 2 mg-N/L: 1.16 (± 0.38)%, 4 mg-N/L: 0.67 (± 0.22)%) with its the abundance significantly higher during condition 1 compared to both conditions 2 and 4 (ANOVA/Tukey, p < 0.05). Further, the abundance of ASV 6 did not change with depth in the ammonia-fed reactor during all conditions whereas the abundance distribution of ASV 4 appeared to be dependent on nitrogen availability (Figure 4A). The abundance of ASV 4 was positively associated with concentrations of ammonia (R=0.344, p < 0.05) and nitrite (R = 0.304, p < 0.05), but the opposite was observed for the abundance of ASV 6 which had a negative association with both ammonia (R=0.367, p < 0.05) and nitrite (R = 0.346, p < 0.05) (Supplemental Figure 4A, B). The relative abundance of *Nitrosomonas-*like ASV 17 remained consistent between conditions (1 mg-N/L: 1.228% (± 0.313), 2 mg-N/L: 1.436% (± 0.393), 4 mg-N/L: 0.969% (± 0.282)) (p < 0.05) (Supplemental Figure 3) with similar relative abundances found in each section of the reactor regardless of nitrogen loading condition (Figure 3B). ASV 54 replaced ASV 17 as the dominant *Nitrosomonas-*like ASV as its average relative increased 27-fold between condition 1 (0.119% (± 0.189)) and 4 (3.233% (± 3.884). The abundance of both ASVs 54 and 62 were positively correlated with the concentration of ammonia (R = 0.503 and R= 0.474, p < 0.05) (Supplemental Figure 4A), indicating their abundance increased in response to higher ammonia concentrations in the ammonia-fed system.

**Figure 4:**
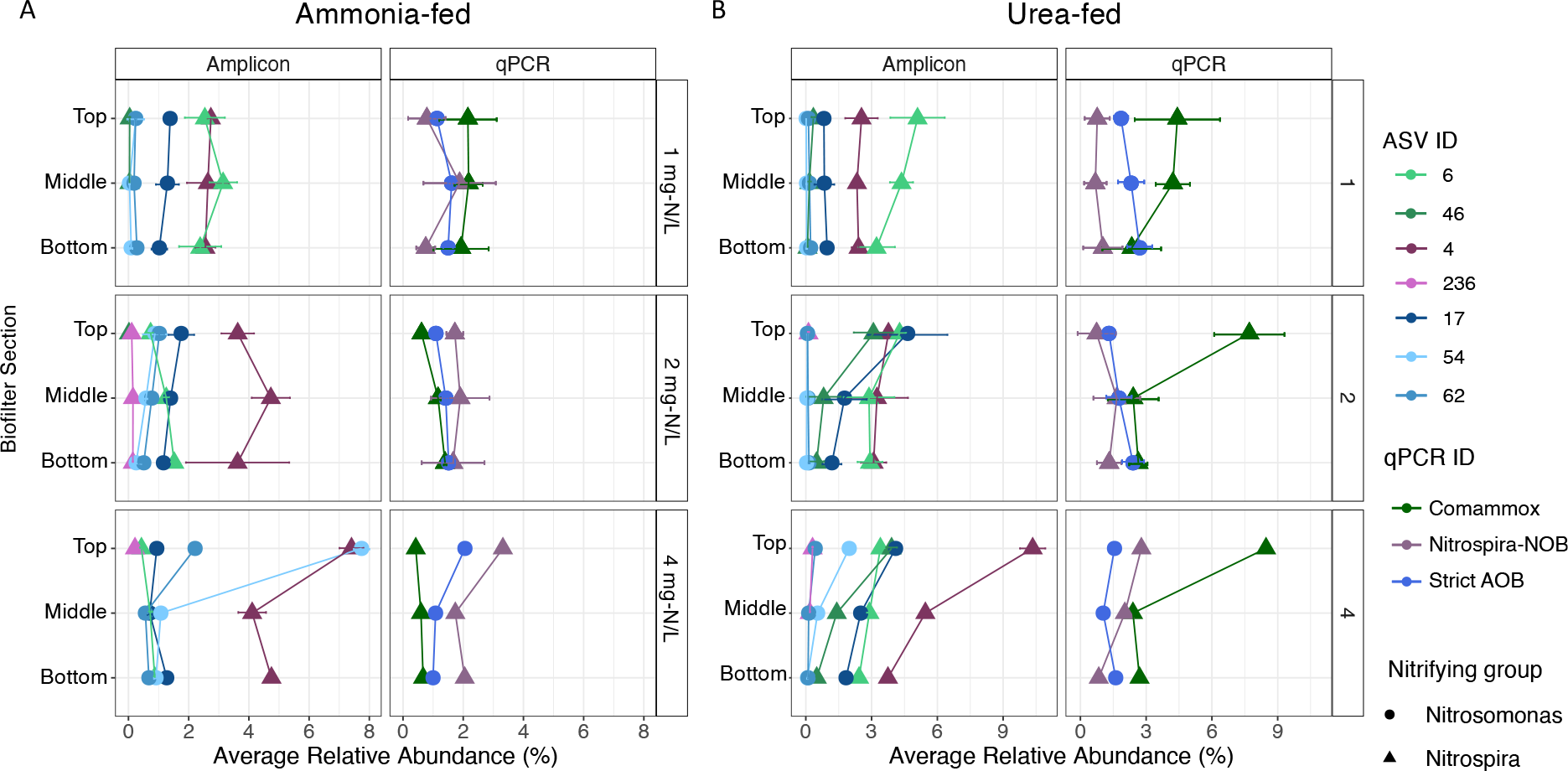
Relative abundance of nitrifier ASVs and qPCR-based relative abundance of comammox bacteria, strict AOB and *Nitrospira-*NOB in the (A) ammonia-fed (A) and (B) urea-fed systems at 1 (top panels), 2 (middle panels), and 4 (bottom panels) mg-N/L. *Nitrosomonas*- and *Nitrospira*-like populations are represented by circles and triangles, respectively. Data points for ASVs are the average relative abundance of nitrifier ASVs calculated in the top, middle and bottom sections of the reactors during each nitrogen loading condition with error bars for standard deviation. Data points for qPCR assays are the average relative abundances of comammox bacteria, strict AOB and *Nitrospira*-NOB in the top, middle and bottom sections of the reactors during each nitrogen loading condition with error bars for standard deviation.

While the abundance of *Nitrosomonas-*like ASV 54 increased 40-fold with increased influent nitrogen concentration in the urea-fed system, ASV 17 remained the dominant ASV with its relative abundance increasing from 0.875% (± 0.310) in condition 1 to 2.450% (± 1.923) in condition 3. In contrast, *Nitrosomonas-*like ASV 62, which increased in abundance proportional to ammonia concentration in the ammonia-fed system, demonstrated no significant change between any of the urea conditions (p > 0.05) and remained at low relative abundance suggesting it was outcompeted in the urea-fed system. While ASVs 4 and 6 were still dominant *Nitrospira*-like ASVs in the urea-fed system, another *Nitrospira*-like ASV (46) increased in abundance from 0.184% (± 0.144) to 1.414% (± 1.361) to 1.934% (± 1.761) for conditions 1, 2, and 3, respectively; ASV 46 was only detected in condition 1 and 2 in the ammonia-fed system at extremely low abundance (<0.006%) thus showing a clear preference for the urea-fed systems (Supplemental Figure 3). It’s abundance also changed with reactor depth during conditions 2 and 4 in the urea-fed system where its abundance was higher in the top section compared to lower portions (Figure 4B). Consistent with the ammonia-fed system, the abundance of *Nitrospira lenta*-like ASV 4 was positively associated with ammonia concentrations (R = 0.289, p < 0.05) (Supplemental Figure 4C) and that of ASV 6 was negatively associated with nitrite concentrations (R= 0.398, p < 0.05) in the urea-fed system (Supplemental Figure 4D).

Comammox bacterial abundance based on qPCR assays was significantly lower in conditions 2 (1.06 (± 0.35) %) and 3 (0.56 (± 0.13) %) compared to its abundance during condition 1 (2.08 (± 0.71)) %) (ANOVA/Tukey, p < 0.05) in the ammonia-fed system (Figure 4A). Additionally, the relative abundance of comammox bacteria displayed minimal change along sections of the ammonia-fed reactor during all conditions which aligns with the trends observed for ASV 6. The relative abundance of *Nitrospira*-NOB assessed by qPCR increased with each nitrogen loading condition (1 mg-N/L: 1.14 (± 0.88) %, 2 mg-N/L: 1.76 (± 0.73) %, 4 mg-N/L: 2.37 (± 0.84) %) though its abundance was not significantly different between each of them (ANOVA/Tukey, p > 0.05). However, its relative abundance was highest overall (3.32%) during condition 3 in the top section where ammonia and nitrite availability was considerably higher compared to the other two nitrogen loading conditions. Thus, *Nitrospira*-NOB likely benefited from increased availability of ammonia and nitrite during higher nitrogen loading conditions whereas comammox bacteria preferred the lowest nitrogen condition with limited ammonia availability in the ammonia-fed system. The overall highest abundance of strict AOB and *Nitrospira*-NOB occurred in the top section of the reactor during condition 3.

qPCR-based abundance of comammox bacteria was significantly higher in all urea-fed conditions compared to its abundance in any of the ammonia-fed conditions (Figure 4A, B) which was also observed for comammox-like ASVs. The combined abundance of comammox-like ASVs was strongly correlated with the qPCR-based abundance of comammox bacteria (Pearson R = 0.92, p < 0.05) (Supplemental Figure 5). Further, qPCR and ASV based abundances agreed that comammox bacteria were the dominant ammonia oxidizers regardless of nitrogen loading condition in the urea-fed system. The qPCR-based abundance of *Nitrospira*-NOB in the urea-fed system was lower than that of comammox bacteria during each condition (Figure 4B). However, increased abundance of comammox bacteria did not result in decreased abundance of *Nitrospira*-NOB (Pearson R = 0.10, p > 0.05) (Figure 5A). This is in contrast to the ammonia-fed system where comammox bacteria and *Nitrospira*-NOB populations demonstrated a significant negative association (Pearson R = -0.48, p < 0.05) (Figure 5B). No significant associations between the abundance of comammox bacteria and strict AOB, and *Nitrospira*-NOB and strict AOB in either the ammonia- or urea-fed systems (Supplemental Figure 6A-D).

**Figure 5:**
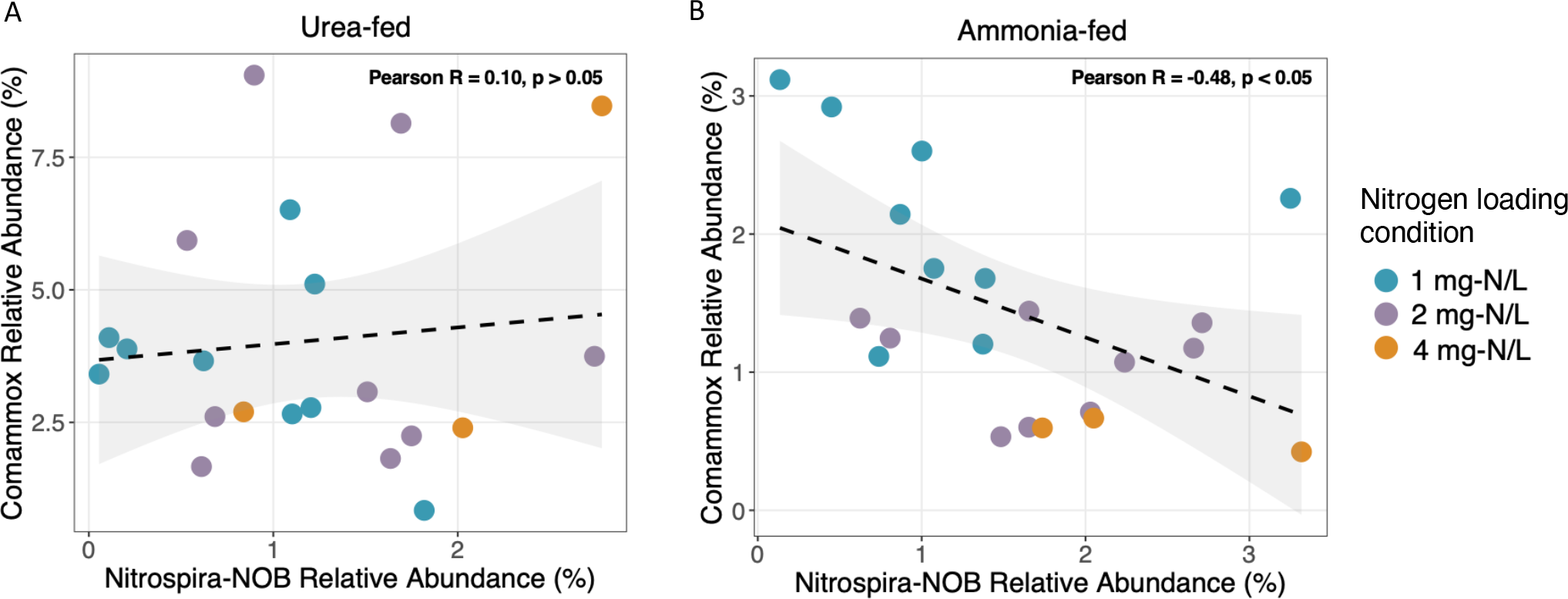
(A) Lack of any relationship between the abundance of comammox bacteria and *Nitrospira*-NOB in the urea-fed system contrasts with (B) significant negative association between the abundance of comammox bacteria and *Nitrospira*-NOB in the ammonia-fed system.

### Phylogeny and metabolism of nitrifier metagenome assembled genomes (MAGs)

200 and 172 MAGs were recovered from the metagenomic assemblies from the ammonia- and urea-fed systems, respectively. The nitrifier community in the ammonia-fed system was comprised of three *Nitrosomonas*-like MAGs and two *Nitrospira*-like MAGs (one classified as Nitrospira_F and one classified as Nitrospira_D) which aligns with the number of dominant nitrifier ASVs in the ammonia-fed system (Table 1). Nitrifier MAGs assembled from the urea-fed system also mirrored the number of dominant nitrifier ASVs with three *Nitrospira*-like (two Nitrospira_F, one Nitrospira_D) and three *Nitrosomonas*-like MAGs. While additional *Nitrosomonas*-like MAG was assembled from the urea-fed GAC sample, it was extremely low quality (completeness < 10%).

**Table 1:** Quality statistics for nitrifier MAGs assembled from GAC taken from the ammonia- and urea-fed reactors.

| MAG name | Classification | Reactor | Completeness (%) | Redundancy (%) |
| --- | --- | --- | --- | --- |
| *Nitrospira_D1_A* | *Nitrospira_D* sp002083555 | Ammonia | 92.27 | 3.91 |
| *Nitrospira_F1_A* | *Nitrospira_F* | Ammonia | 88.88 | 3.69 |
| *Nitrosomonas_1_A* | *Nitrosomonas* | Ammonia | 80.3 | 0.48 |
| *Nitrosomonas_2_A* | *Nitrosomonas* sp016708955 | Ammonia | 97.38 | 0.51 |
| *Nitrosomonas_3_A* | *Nitrosomonas* | Ammonia | 92.34 | 0.03 |
| *Nitrospira_F1_U* | *Nitrospira_F* | Urea | 92.11 | 71.93 |
| *Nitrospira_D1_U* | *Nitrospira_D* sp002083555 | Urea | 94.09 | 4.82 |
| *Nitrospira_F2_U* | *Nitrospira_F* sp002083565 | Urea | 84.65 | 6.41 |
| *Nitrosomonas_1_U* | *Nitrosomonas* | Urea | 98.72 | 0.48 |
| *Nitrosomonas_2_U* | *Nitrosomonas* sp016708955 | Urea | 93.38 | 0.51 |
| *Nitrosomonas_3_U* | *Nitrosomonas* | Urea | 93.54 | 0.03 |
| *Nitrosomonas_4_U* | *Nitrosomonas* | Urea | 6.22 | 0 |

Phylogenetic analysis with 91 single copy core genes clustered Nitrospira_F1_A with clade A comammox bacteria (Figure 6A) and it showed high sequence similarity (∼94% ANI) with Nitrospira sp Ga0074138 which is a comammox bacteria MAG previously assembled by Pinto et al. (2015) from GAC obtained from the same reactor. Nitrospira_F1_A shared extremely high sequence similarity (> 99% ANI) with Nitrospira_F1_U assembled from the urea-fed system, suggesting the two MAGs were likely the same population (Supplemental Figure 7). Another *Nitrospira* MAG (Nitrospira_F2_U) assembled from the urea-fed system sample was placed within comammox clade A, but clustered separately with other drinking water related comammox MAGs (Nitrospira sp. ST-bin4 and SG-bin2). This MAG shared less than 80% ANI with all other *Nitrospira* MAGs in this study. Phylogenetic placement of hydroxylamine oxidoreductase (HAO) gene sequences present in all comammox MAGs in this study were grouped into clade A2 comammox bacteria (data not shown). The remaining two *Nitrospira* MAGs (Nitrospira_D1_A and Nitrospira_D1_U) clustered with *Nitrospira*-NOB belonging to lineage II (Figure 6A) with *Nitrospira lenta* and other Nitrospira-NOB obtained from a drinking water system (Nitrospira sp. ST-bin5) and rapid sand filter (RSF 13 and CG24D). Nitrospira_D1_A and Nitrospira_D1_U from this study shared over 99% sequence similarity indicating they are the same population (Supplemental Figure 7).

**Figure 6:**
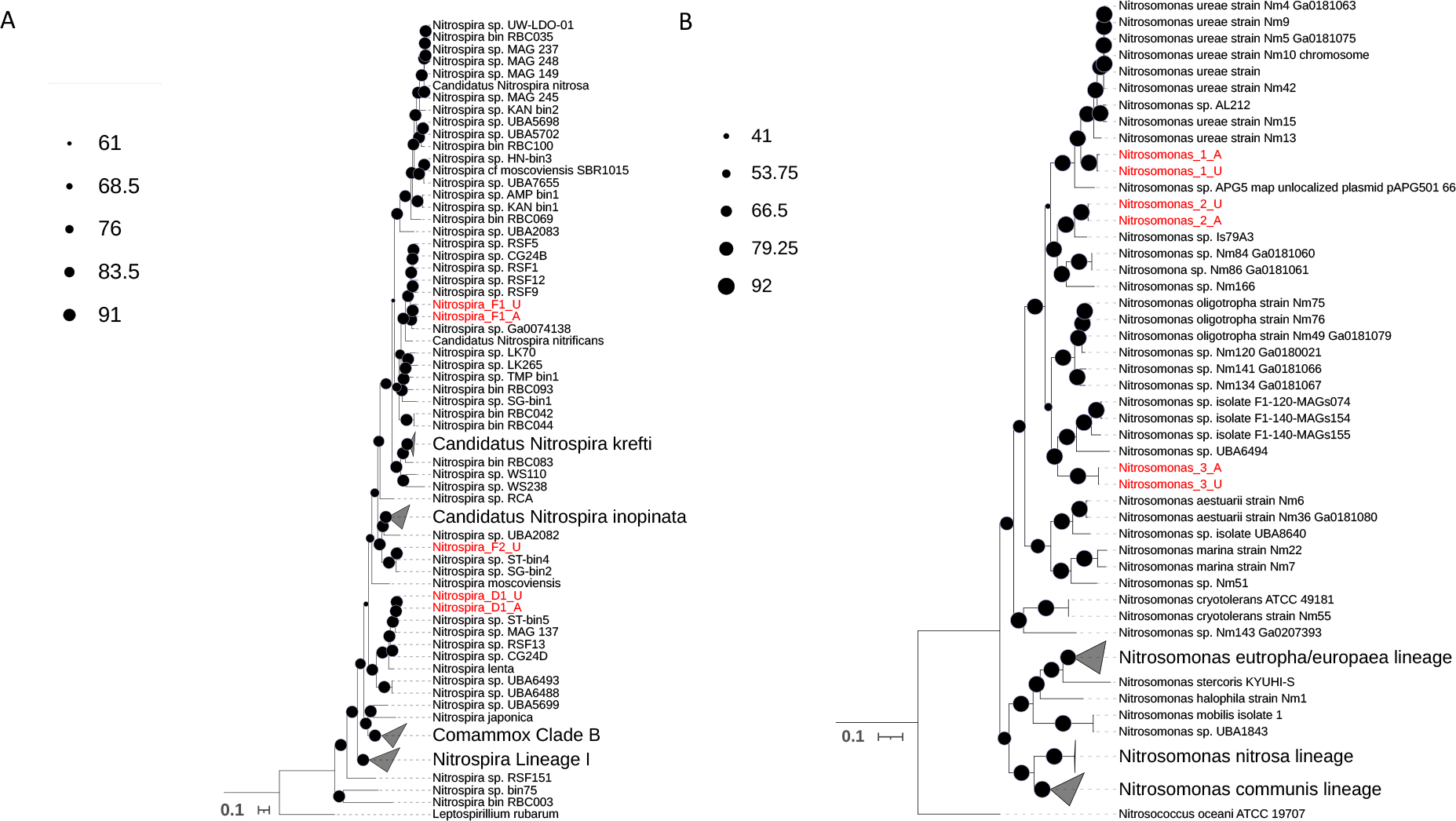
Maximum likelihood trees for (A) Nitrospira and (B) Nitrosomonas based on a set of bacterial single copy core genes. MAGs assembled in this study are labelled in red and reference genomes and MAGs are black.

ANI comparisons between the *Nitrosomonas*-like MAGs assembled from the ammonia- (Nitrosomonas_1_A, Nitrosomonas_2_A, Nitrosomonas_3_A) and urea-fed (Nitrosomonas_1_U, Nitrosomonas_2_U, Nitrosomonas_3_U) systems revealed the same set of three *Nitrosomonas*-like MAGs were assembled from both samples (Supplemental Figure 8). Within sample ANI comparisons showed that the three *Nitrosomonas*-like MAGs shared less than 95% ANI suggesting they were separate species. Phylogenomic placement of Nitrosomonas MAGs in this study affiliated them with *Nitrosomonas* cluster 6a which are known for their oligotrophic physiologies (Koops 2001). Nitrosomonas_1_A and Nitrosomonas_1_U (ANI > 99%) clustered with *Nitrosomonas ureae* while Nitrosomonas_2_A and Nitrosomonas_2_U (ANI > 99%) grouped with Nitrosomonas Is79A3 (Figure 6B). Nitrosomonas_3_A and Nitrosomonas_3_U (ANI > 99%) clustered with uncultured Nitrosomonas MAGs that were still within the cluster 6a grouping. Out of all reference comparisons, Nitrosomonas MAGs from this study shared the highest similarity to Nitrosomonas ureae strain Nm5 Ga0181075 101 (ANI = 83%, Nitrosomonas_1_A and Nitrosomonas_1_U), Nitrosomonas Is79 (ANI = 89%, Nitrosomonas_2_A and Nitrosomonas_2_U), and Nitrosomonas sp. Nm141 Ga0181066 101 (ANI = 79%, Nitrosomonas_3_A and Nitrosomonas_3_U).

Dereplication of MAGs from urea- and ammonia-fed systems resulted in three Nitrosomonas-like MAGs, one *Nitrospir*a-NOB MAG and two comammox bacteria-like MAGs. Filtered reads from the ammonia-fed system were mapped to the set of dereplicated MAGs, revealing all nitrifier MAGs had 99% breath of coverage (i.e., percent of genome covered by reads) in both systems. However, the comammox MAG that was assembled only from the urea-fed sample (Nitrospira_F2_U) had very low relative abundance (0.065%) in the ammonia-fed system which could explain why it was not assembled. Comparably, relative abundances of comammox bacteria MAGs, Nitrospira_F1_A/Nitrospira_F1_U and Nitrospira_F2_U, were approximately 12 and 5-fold higher in the urea-fed system (∼7.81% Nitrospira_F1_A/Nitrospira_F1_U, 0.30% Nitrospira_F2_U) than in the ammonia-fed system (∼0.66 Nitrospira_F1_A/ Nitrospira_F1_U, 0.065% Nitrospira_F2_U). These results align with both the qPCR-based abundance of comammox bacteria and abundance of comammox-like ASVs 6 and 46 in the urea-fed system being substantially higher than their abundance in the ammonia-fed system. Thus, based on abundance trends of the comammox-like ASVs 6 and 46 and comammox MAGs, we associate ASV 6 with the comammox bacterial population belonging to Nitrospira_F1_A/ Nitrospira_F1_U while ASV 46 is associated with Nitrospira_F2_U.

Similar to our previous study, comammox MAGs (Nitrospira_F1_A, Nitrospira_F1_U, Nitrospira_F2_U) contained genes for urea degradation (*ureCAB*) and transportation (*urtACBCDE*). Strict AOB MAGs Nitrosomonas_1_A and Nitrosomonas_1_U also possessed these genes for ureolytic activity which aligns with their sequence similarity to and clustering with *Nitrosomonas ureae*. The other Nitrosomonas MAGs (Nitrosomonas_2_A, Nitrosomonas_2_U, Nitrosomonas_3_A, Nitrosomonas_3_U) only encoded a single urea accessory gene (*ureJ*) and gene encoding for urea carboxylase. An unbinned *Nitrosomonas*-associated *ureC* gene was found in the metagenome assembly from the urea-fed system suggesting another urease-positive strict AOB MAGs could have been present in the system. *Nitrospira* MAGs (Nitrospira_D1_A and Nitrospira_D1_U) did not contain urease genes; however, unbinned genes for *ureA* with 100% sequence ID match to strict NOB *Nitrospira lenta* were detected in the metagenome assembly for both samples, suggesting Nitrospira-NOB were urease-positive.

## DISCUSSION

### Nitrite accumulation in ammonia-fed but not urea-fed system may be associated with NOB inhibition and with the rate of ammonia production from urea

Strict AOB and *Nitrospira*-NOB were the dominant nitrifiers in the ammonia fed systems and particularly at higher ammonia concentrations with the ammonia oxidation rates being consistently higher than the nitrite oxidation rates leading to nitrite accumulation. While nitrite accumulation occurred in the ammonia-fed reactor for condition 3, *Nitrospira*-NOB were more abundant than both AOB and comammox bacteria. It could be possible that despite their high abundance, *Nitrospira*-NOB were impacted by higher ammonia concentrations of conditions 2 and 3. Fujitani et al. (2020) observed that the average K_m_ value for nitrite (0.037 mg/L) attributed to a *Nitrospira*-NOB strain originating from a drinking water treatment plant increased five-fold to approximately 0.18 mg-N/L NO_2-_ in the presence of free ammonia concentrations around 0.85 mg NH_3_-N/L. Thus, decreased nitrite affinity could have impacted the ability of this *Nitrospira* strain to oxidize low nitrite concentrations depending on the concentration of free ammonia. Further, in wastewater systems, suppression of strict NOB activity can be achieved at ammonia concentrations were higher than 5 mg-N/L (Poot et al., 2016). Here in the ammonia-fed system, average ammonia concentrations observed in the top section of the reactor during condition 2 (0.89 mg NH_3_/L) and 4 (2.64 mg NH_3_/L) were in line with free ammonia concentrations shown to impact nitrite affinity of *Nitrospira*-NOB strain KM1 in Fujitani et al. (2020), thus explaining nitrite accumulation. In the urea-fed system, urease-positive nitrifiers, including comammox bacteria and *Nitrospira*-NOB, regulated ammonia production and thus potentially controlled ammonia availability. While ammonia did accumulate during the highest nitrogen loading condition in the urea-fed reactor, however, unlike the ammonia-fed reactors, comammox bacterial abundance did not decrease and there was no nitrite accumulation. This is likely because the highest ammonia concentrations in the urea-fed reactor were consistently lower than the highest concentrations in the ammonia-fed reactors and thus comammox bacteria were not outcompeted by AOB and both comammox and *Nitrospira*-NOB were likely not inhibited.

### Increased ammonia availability in ammonia-fed reactor detrimentally impacted comammox bacterial populations

Consistent with reported higher ammonia affinity (i.e., lower Km(app)) of comammox bacteria compared to strict AOB (Kits 2017, Sakoula 2020, Ghimire-Kafle 2023), comammox bacteria did indeed dominate over AOB only during the lowest nitrogen loading condition in the ammonia-fed system. Though strict AOB were affiliated with *Nitrosomonas* cluster 6a characterized with higher ammonia affinities (Km(app)=0.24-3.6 μM (Koops et al., 2006)) compared to other AOB, the reported ammonia affinity for comammox bacteria is still substantially higher for comammox bacteria (Km(app)=63 nM). In addition to ammonia affinity, it is very likely that ammonia tolerance played a role as partial inhibition of ammonia oxidation activity by *Ca* Nitrospira kreftii has been reported at ammonia concentrations as low as 0.425 mg/L which is within the range of ammonia concentrations observed in the ammonia-fed reactor (0.25-2 mg-N/L) during conditions 2 and 3. However, ammonia sensitivity resulting in partial inhibition of ammonia oxidation has not been observed for *Ca* Nitrospira inopinata (Kits 2017), and *Ca* Nitrospira nitrosa-like comammox bacteria in a wastewater systems were ammonia concentrations are higher (Cotto et al., 2020, 2023; K. Vilardi et al., 2023). Comammox bacteria in this study may be adapted to low ammonia concentrations and were most similar to other clade A2 comammox bacteria obtained from low ammonia environments. Thus, continuous exposure to elevated ammonia concentrations could be responsible for the observed reduction in abundance of comammox bacteria via inhibition.

### Increase in urea concentration favored *Nitrospira* bacteria including comammox bacteria

Comammox bacteria were the dominant nitrifier across all conditions in the in the urea-fed system with its overall abundance significantly higher in the urea-fed system compared to its abundance in the ammonia-fed system. Urea preference of comammox bacteria was also supported by the highest abundances of comammox bacteria consistently observed in the top of the urea-fed reactor where urea was most available and the emergence of a second low abundance comammox population in the urea-fed reactor. Our observation is similar to other reports of enrichment of very different comammox populations at much higher urea concentrations (J. Li et al., 2021; Zhao et al., 2021). Though we are unable to identify the exact reason for comammox bacterial preference for growth on urea, it could be a combination of metabolic traits associated with urea uptake and utilization. Specifically, comammox bacteria may balance the rate of ammonia production from urea with its ammonia oxidation rate, thus maximizing ammonia availability while also maintaining ammonia concentrations at non-inhibitory levels. Further, additional urea transporters are found in comammox genomes that are absent in other *Nitrospira* including an outer-membrane porin (*fmdC*) for uptake of short-chain amides and urea at low concentrations and a urea carboxylase-related transport (*uctT*) (Palomo 2018). Thus, the enhanced ability to uptake urea and regulate its conversion to ammonia balanced with its ammonia oxidation rates may underpin comammox bacterial preference for urea. Estimating the kinetic parameters such as comammox bacteria’s affinity for urea and uptake rate relative to other nitrifiers and ammonia production relative to its own ammonia oxidation rates would be extremely useful for assessing their overall preference for urea.

### Nitrogen source drives potential competitive and co-operative dynamics between aerobic nitrifiers

In these continuous flow reactors and our previous batch microcosm experiments (K. J. Vilardi et al., 2022), we observed nitrogen source-dependent dynamics between the abundance of comammox bacteria and *Nitrospira*-NOB. We hypothesized that tight metabolic coupling exists between strict AOB and *Nitrospira*-NOB when urea is supplied due to reciprocal feeding mediated by the two groups. Here, the production of nitrite can be controlled by urease-positive *Nitrospira*-NOB via cross feeding ammonia to strict AOB, who in turn provide nitrite at a rate at which *Nitrospira*-NOB can consume it. This dynamic between canonical nitrifiers substantially contrasts with their relationship when only ammonia is provided as *Nitrospira*-NOB are fully dependent on strict AOB to provide them nitrite. Therefore, a negative relationship between comammox and canonical NOB *Nitrospira* when only ammonia is available may reflect comammox bacteria limiting *Nitrospira*-NOB access to nitrite (produced by AOB) by performing complete ammonia oxidation to nitrate. In contrast, at high ammonia concentration, comammox bacteria may in fact be a source of nitrite for *Nitrospira*-NOB as their ammonia oxidation rates are faster than their nitrite oxidation rates, and their affinities for nitrite are lower than that of *Nitrospira*-NOB (Daims et al., 2015; van Kessel et al., 2015). Supplementation with urea eliminates this potential comammox-NOB negative association as both nitrifiers are urease-positive and potentially produce ammonia themselves for different purposes (i.e., comammox produce their own ammonia, strict NOB provide ammonia to strict AOB). Competition for urea would then be determined by the urea affinity and uptake rates which are currently unknown. However, in this study, we show that increased abundance of comammox bacteria did not result in decreased abundance of *Nitrospira-*NOB in the urea-fed system. This suggests that the apparent competitive dynamics between these nitrifiers is reduced when an alternative nitrogen source is available compared to ammonia which induced a competitive relationship.

In this study, the impact of nitrogen source and availability on nitrifying communities was evaluated in continuous flow column reactors supplied either ammonia or urea and operated over three different nitrogen loading conditions. Consistent with our previous batch microcosm experiments (Vilardi et al, 2022), we show that different nitrogen sources and loadings distinctly shape the nitrifying community. Direct supply of ammonia favored a combination of AOB and NOB particularly as the nitrogen loadings were increased, with decrease in comammox bacterial abundance was likely associated with ammonia-based inhibition. Ammonia availability has been considered an important niche differentiating factor between comammox bacteria and strict AOB, and here we show it may also be a significant factor for shaping populations of comammox bacteria. In contrast, the urea provision promoted the abundance of multiple comammox populations along with strict AOB and *Nitrospira*-NOB. With urea as a nitrogen source, nitrification can be initiated by urease-positive nitrifiers controlling ammonia production and its availability which in turn significantly impacted nitrification process performance.

## Funding sources

This work was supported by NSF Graduate Research Fellowship and Cochrane Fellowship to KV and by NSF Award number: 2203731.

## Notes

### Competing Interest Statement

The authors have declared no competing interest.

