## Supplemental materials for "Comammox bacterial preference for urea influences its interactions with aerobic nitrifiers"

**Supplemental Table 1**: Sequences of primers, PCR reaction and thermocycling conditions, and qPCR efficiencies for quantification of comammox bacteria *amoB*, *Nitrospira* 16S rRNA, AOB 16S rRNA and total bacteria 16S rRNA.

| **Target** | **Primer Set** | **Forward Primer sequence** | **Reverse Primer sequence** | **Final concentration in PCR reaction** | **Annealing Temp (°C) /Time (sec)** | **Amplicon length (bp)** | **Reference** |
|---|---|---|---|---|---|---|---|
| Comammox clade A amoB gene | Mod_CMX_amoB 148F/485R | TGGTAYGAYACNSARTGGG | CCNGTGATRTCCATCCA | 0.5 mM F/R | 52/45 | 337 | (1) |
| Nitrospira 16S rRNA gene | Nspra675F - 746R | GCGGTGAAATGCGTAGAKATCG | TCAGCGTCAGRWAYGTTCCAGAG | 0.5 mM F/R | 58/30 | 93 | (2) |
| AOB 16S rRNA gene | CTO189FA/B/C-RT1R | GGAGRAAAGCAGGGGATCG+ GGAGGAAAGTAGGGGATCG | CGTCCTCTCAGACCARCTACTG | 0.4 mM F/R | 57/30 | 116 | (3) |
| Total Bacteria 16S rRNA gene | F515 - R806 | GTGCCAGCMGCCGCGGTAA | GGACTACHVGGGTWTCTAAT | 0.2 mM F/0.4 mM R | 50/15 | 291 | (4) |


1. Vilardi KJ, Cotto I, Sevillano M, Dai Z, Anderson CL, Pinto A. 2022. Comammox *Nitrospira* bacteria outnumber canonical nitrifiers irrespective of electron donor mode and availability in biofiltration systems. FEMS Microbiology Ecology 98.
2. Graham DW, Knapp CW, Van Vleck ES, Bloor K, Lane TB, Graham CE. 2007. Experimental demonstration of chaotic instability in biological nitrification. ISME J 1:385–393.
3. Hermansson A, Lindgren P-E. 2001. Quantification of Ammonia-Oxidizing Bacteria in Arable Soil by Real-Time PCR. Applied Environmental Microbiology67:972–976.
4. Caporaso JG, Lauber CL, Walters WA, Berg-Lyons D, Lozupone CA, Turnbaugh PJ, Fierer N, Knight R. 2011. Global patterns of 16S rRNA diversity at a depth of millions of sequences per sample. Proceedings of the National Academy of Sciences 108:4516–4522.

**Supplemental Table 2**: Details of gBlock qPCR standards and dilution range for 7-point standard curve (1 order of magnitude difference between points).

| **Target gene** | **Sequence source** | **Gblock sequence** | **Standard curve range (copies/5 µl)** |
|---|---|---|---|
| Comammox clade A amoB | Nitrospira inopinata | CACCGTGAATTGGTATGACACTGAATGGGTGGGGAAAAGC<br>ACTGCGGTAAATGATGTTACATACATGAGGGGCAAGTTTCA<br>TCTGTCTGAAGACTGGCCTCGTGCGGTAGTGAAACCCCAT<br>CGAACGTTCGTCAATGTCGGCTCTCCTAGCTCCGTCTTTGT<br>GCGGTTAAGCACGAAGGTTGGTGGGGTGCCGATGTTTGT<br>GTCTGGTCCTATGGAAATCGGGCGTGATTATGAATATGAG<br>ATCACGTTGAAGGCGAGACTTCCTGGACATCATCACATTCA<br>CCCTATGTTTTCTGTTAAAGAGGCTGGTCCCATTGCCGGA<br>CCGGGTGGGTGGATGGATATCACGGGCCGATACGCT | $10^2$-$10^8$ |
| Nitrospira 16S rRNA/Total bacteria 16S rRNA | Nitrospira inopinata | GGCTAACTTCGTGCCAGCAGCCGCGGTAATACGAAGGTG<br>GCAAGCGTTGTTCGGATTTACTGGGCGTACAGGGAGCGTA<br>GGCGGTTGGGTAAGCCCTCCGTGAAATCTCCGGGCCTAA<br>CCCGGAAAGTGCGGAGGGGACTGCTTGGCTAGAGGATGG<br>GAGAGGAGCGCGGAATTCCCGGTGTAGCGGTGAAATGCG<br>TAGAGATCGGGAGGAAGGCCGGTGGCGAAGGCGGCGCT<br>CTGGAACATTTCTGACGCTGAGGCTCGAAAGCGTGGGGA<br>GCAAACAGGATTAGATACCCTGGTAGTCCACGCCCTAAA | $10^2$-$10^8$ (Nitrospira) $10^3$-$10^9$ (Total Bacteria) |
| AOB 16S rRNA gene | Nitrosomonas europaea | CATATCTCTGAGGAGAAAAGCAGGGGATCGCAAGACCTTG<br>CGCTAAAGGAGCGGCCGATGTCTGATTAGCTAGTTGGTGG<br>GGTAAAGGCTTACCAAGGCAACGATCAGTAGTTGGTCTGA<br>GAGGACGGCCAACCACA | $10^2$-$10^8$ |

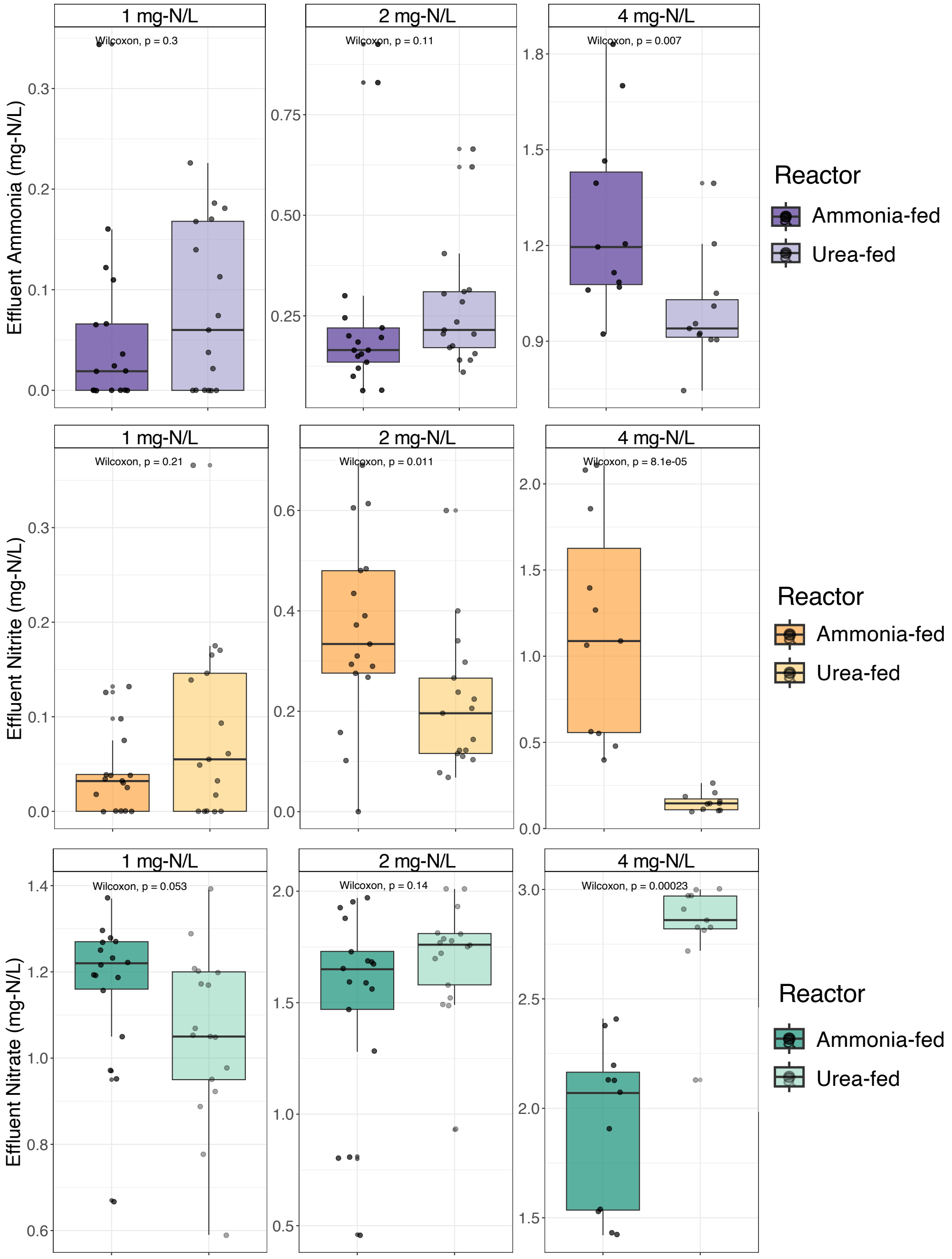


**Supplementary Figure 1**: Effluent concentrations of ammonia (purple), nitrite (orange), and nitrate (green) measured twice per week in the ammonia- and urea-fed biofiltration systems during all nitrogen loading conditions. Facet labels indicate input inorganic nitrogen concentration.

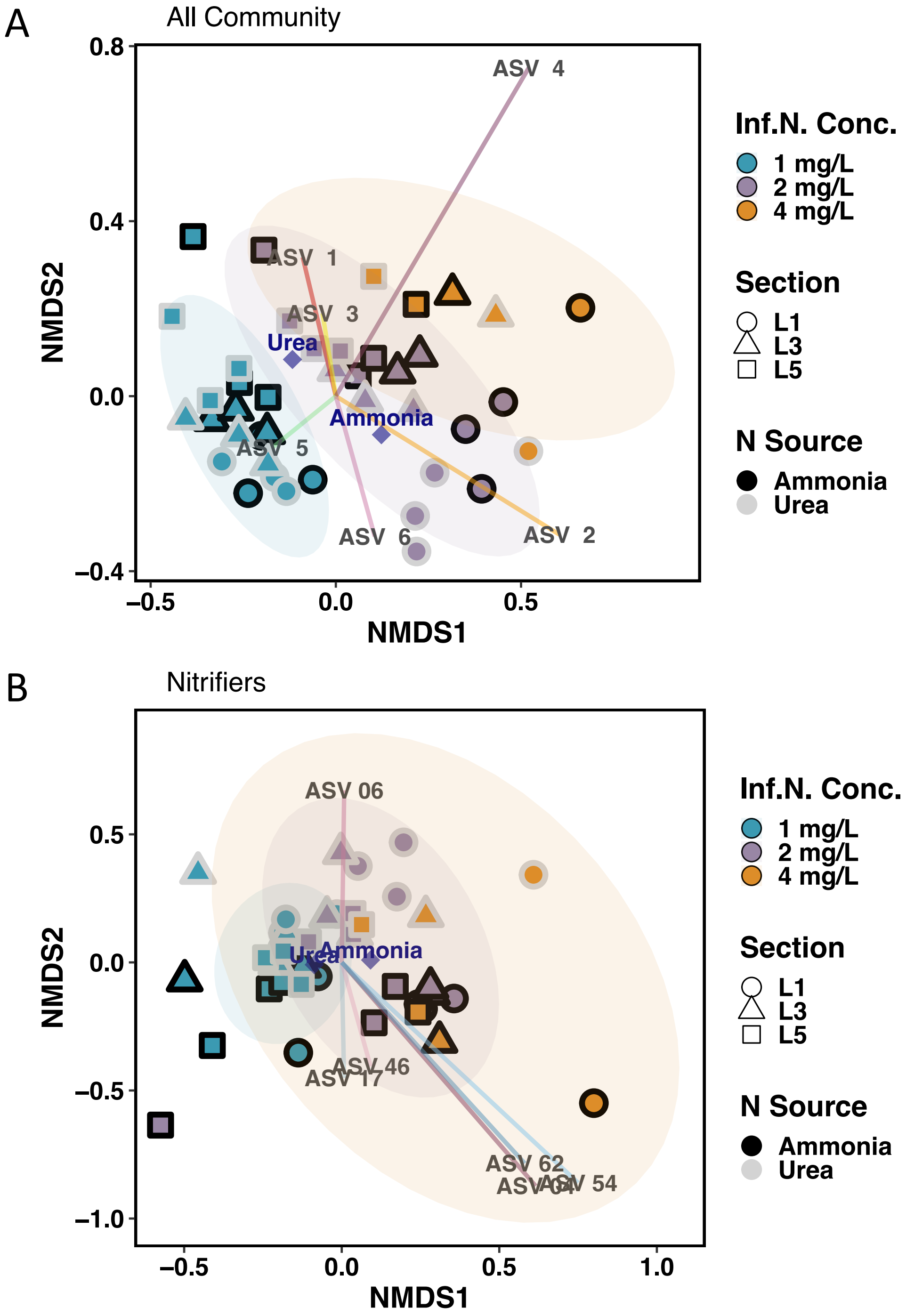


**Supplementary Figure 2:** (A) NMDS plots constructed with the abundance tables of all ASVs, (B) all nitrifier ASVs. Blue-, purple- and orange-colored points are GAC samples collected during conditions 1, 2, and 3, respectively. Shape symbolizes the reactor depth the GAC samples were taken from (L1 = top (circle), L3 = middle (triangle), and L5 = bottom (square)). The outline color of shapes represents the system the GAC was collected from (gray = urea-fed, black = ammonia-fed).

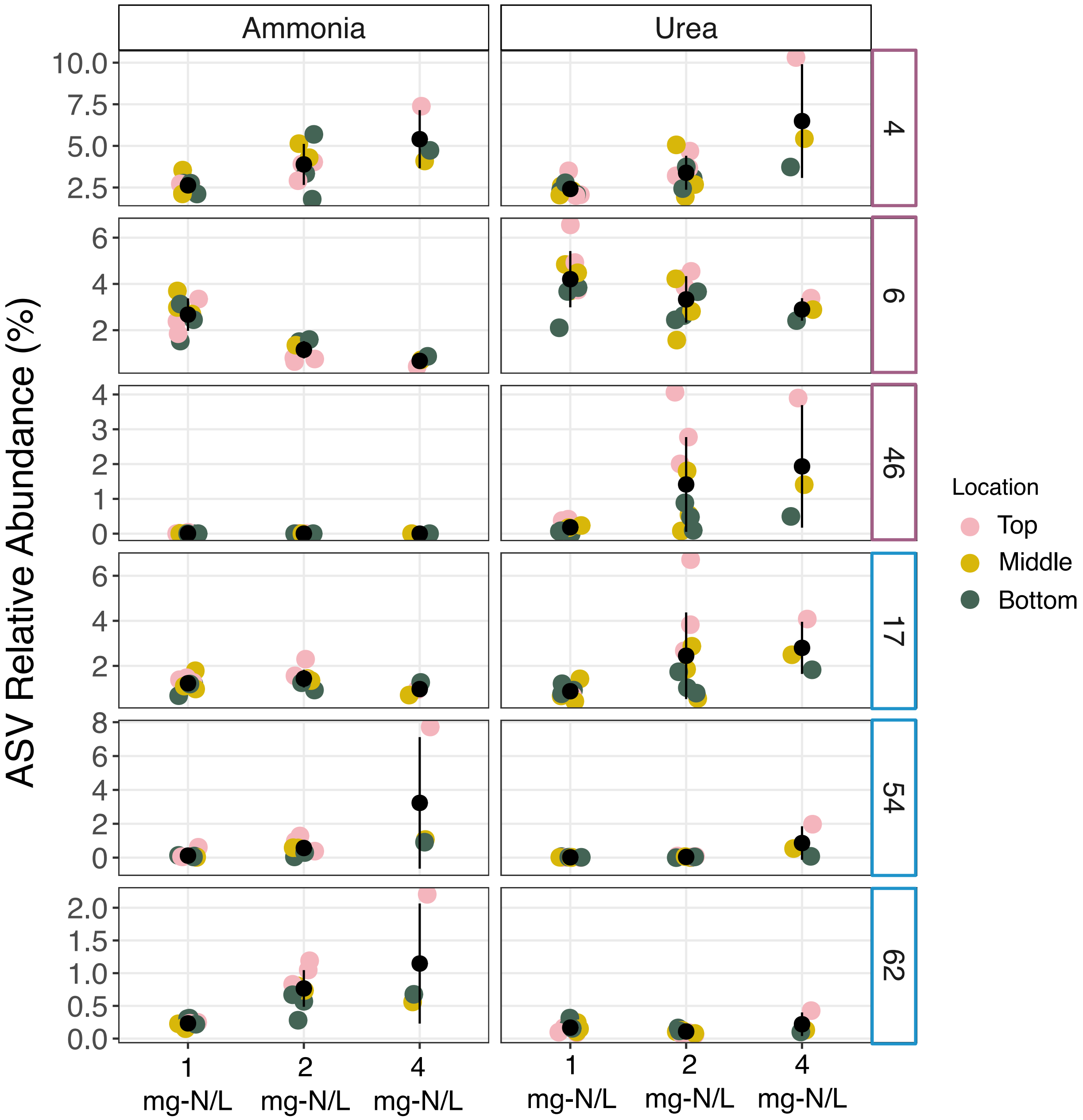


**Supplementary Figure 3**: Average relative abundance of nitrifier ASVs in the ammonia- and urea-fed biofiltration systems during each condition (black data points). Each colored data point is an average of technical replicates. *Nitrospira* and *Nitrosomonas* ASVs are outlined in purple and blue, respectively.

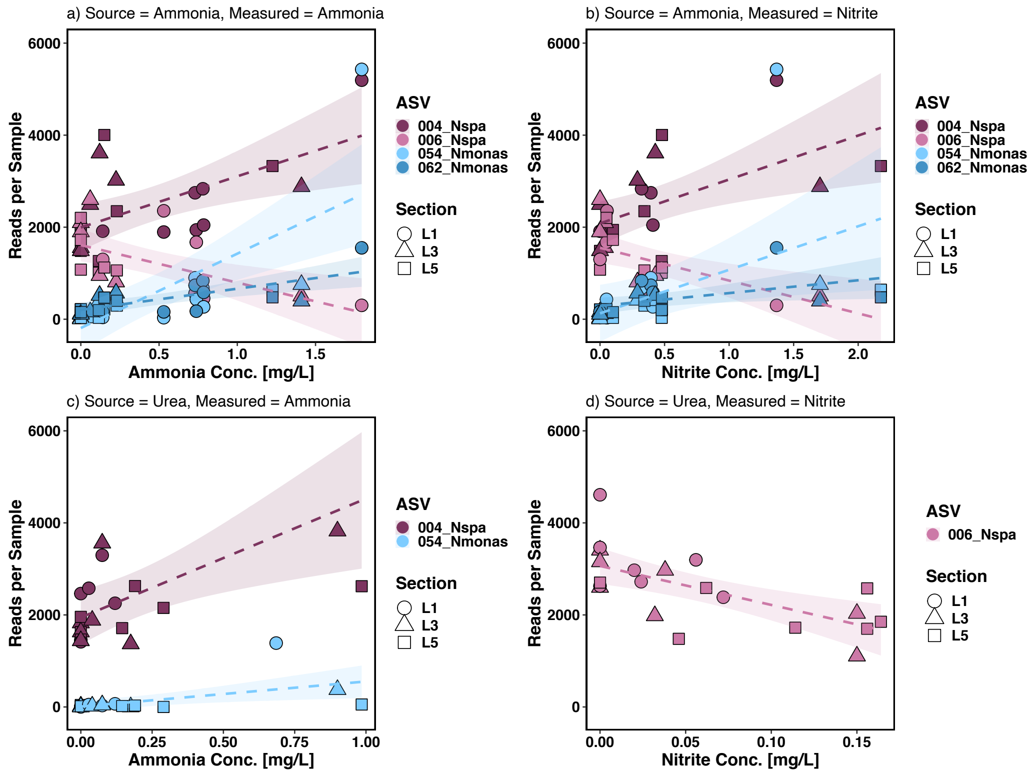


**Supplemental Figure 4**: Significant correlations between the abundance of nitrifier ASVs and concentrations of ammonia (A) and nitrite (B) measured in the ammonia-fed system and concentrations of ammonia (C) and nitrite (D) measured in the urea-fed system.

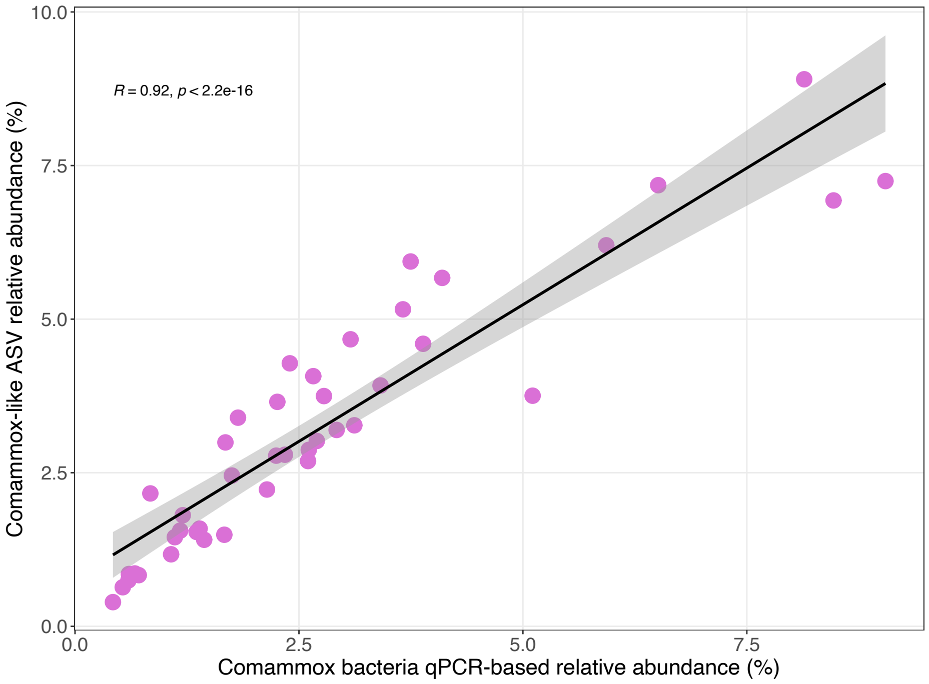


**Supplemental figure 5**: Significant positive correlation between the abundance of comammox bacteria assessed with qPCR and amplicon sequencing detected using Pearson correlation.

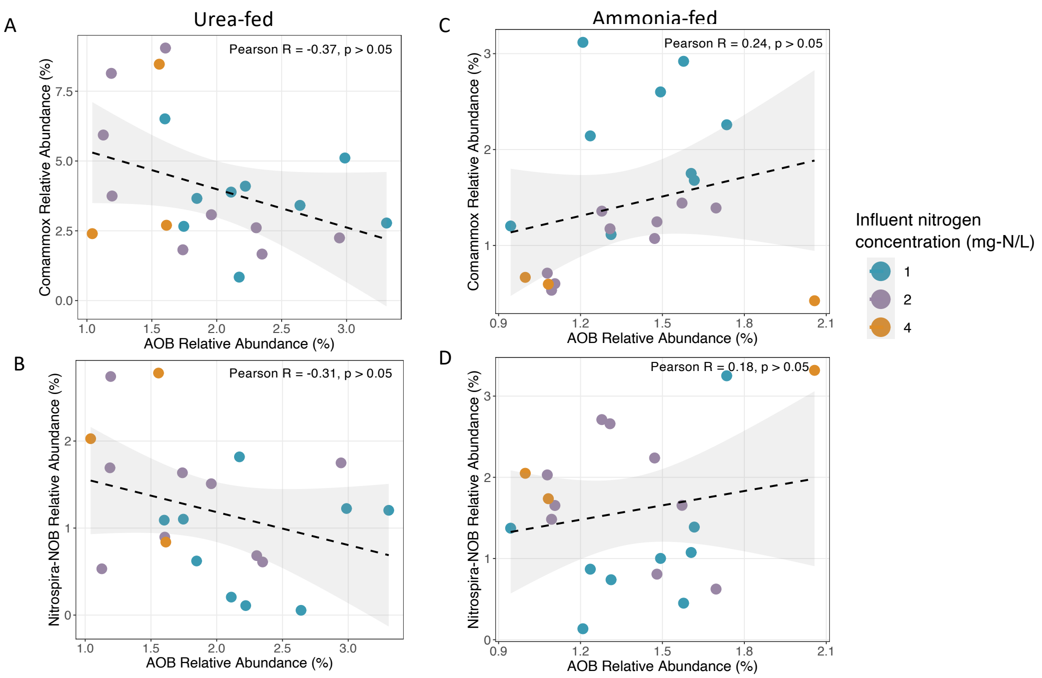


**Supplementary Figure 6**: Lack of association between the qPCR-based abundance of comammox bacteria and strict AOB (A and C), and *Nitrospira*-NOB and strict AOB (B and D) in the ammonia- and urea-fed systems.

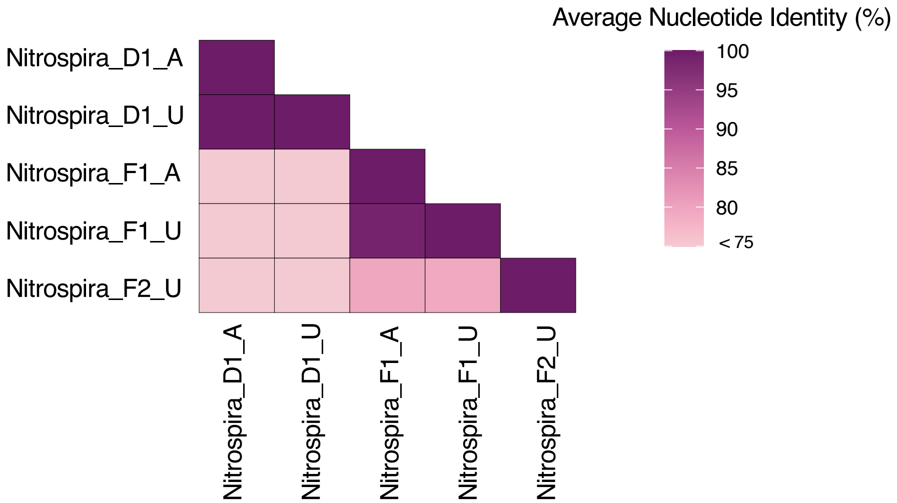


**Supplemental Figure 7**: Average nucleotide identities of Nitrospira MAGs recovered in this study.

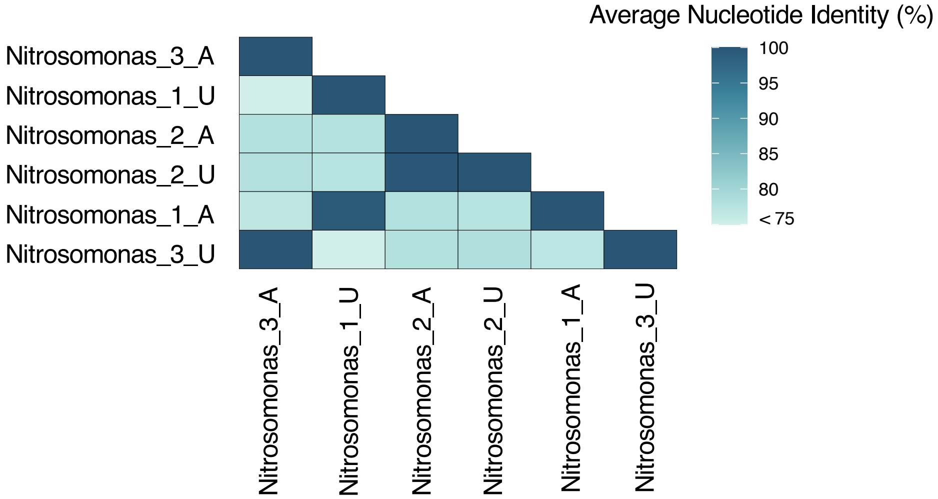


**Supplemental Figure 8**: Average nucleotide identities of Nitrosomonas MAGs recovered in this study.
